# The impact of tumor stromal architecture on therapy response and clinical progression

**DOI:** 10.1101/451047

**Authors:** Philipp M. Altrock, Nara Yoon, Joshua A. Bull, Hao Wu, Javier Ruiz-Ramírez, Daria Miroshnychenko, Gregory J. Kimmel, Eunjung Kim, Robert J. Vander Velde, Katarzyna Rejniak, Brandon J. Manley, Fabian Spill, Andriy Marusyk

## Abstract

— Advances in molecular oncology research culminated in the development of targeted therapies that act on defined molecular targets either on tumor cells directly (such as inhibitors of oncogenic kinases), or indirectly by targeting the tumor microenvironment (such as anti-angiogenesis drugs). These therapies can induce strong clinical responses, when properly matched to patients. Unfortunately, most targeted therapies ultimately fail as tumors evolve resistance. Tumors consist not only of neoplastic cells, but also of stroma, whereby “stroma” is the umbrella term for non-tumor cells and extracellular matrix (ECM) within the tumor microenvironment, possibly excluding immune cells^1^. We know that tumor stroma is an important player in the development of resistance. We also know that stromal architecture is spatially complex, differs from patient to patient and changes with therapy. However, to this date we do not understand the link between spatial and temporal changes in stromal architecture and response of tumors to therapy, in space and time. In this project we sought to address this gap of knowledge using a combination of mathematical and statistical modeling, experimental *in vivo* studies, and analysis of clinical samples in therapies that target tumor cells directly (in lung and breast cancers) and indirectly (in kidney cancer). This knowledge will inform therapy choices and offer new angles for therapeutic interventions. Our main question is: how does spatial architecture of stroma impact the emergence or evolution of resistance to targeted therapies, and how can we use this knowledge clinically?

## I. Introduction

Tumors are complex, abnormal tissues, comprised of nests of tumor cells surrounded by stroma. The stroma is the connective tissue, composed of cancer-associated fibroblasts (CAFs), extracellular matrix (produced primarily by CAFs), vasculature, lymphatics and immune cells. A growing body of pre-clinical studies indicates that stroma in general, and CAFs in particular, can have profound impacts on tumor growth, progression and therapy responses^2–4^. Physical barriers imposed by stroma can restrain tumor growth and progression. Stroma provides therapy protection in a wide range of targeted therapies^5,6^. This protection is mediated by paracrine pro-survival signals acting over short distance; therefore, spatial patterns of stroma and cancer cell localization are expected to have a profound impact on stromal protection. Yet, the specific architecture of the stroma is generally ignored during clinical diagnosis and therapy decision-making. The major reason for this omission is that we lack approaches to extract meaning from the relevant data, despite almost universal availability of data on stromal content and spatial CAF-distribution in diagnostic clinical samples. The situation is further complicated by the fact that stroma is spatially complex and heterogeneous and dynamically changing during disease progression and therapy response. This lack of attention to tumor stroma is also fully applicable to clinical diagnostics and decision-making in clear cell renal cell carcinoma (ccRCC), an aggressive epithelial malignancy. Despite availability of targeted therapy directed against neoplastic cells directly (mTOR inhibitors) or indirectly (anti-angiogenic agents), late stage ccRCC remains incurable^7,8^. The different stages of this disease show distinct stromal architectures (Figure 1).

**Figure 1.**
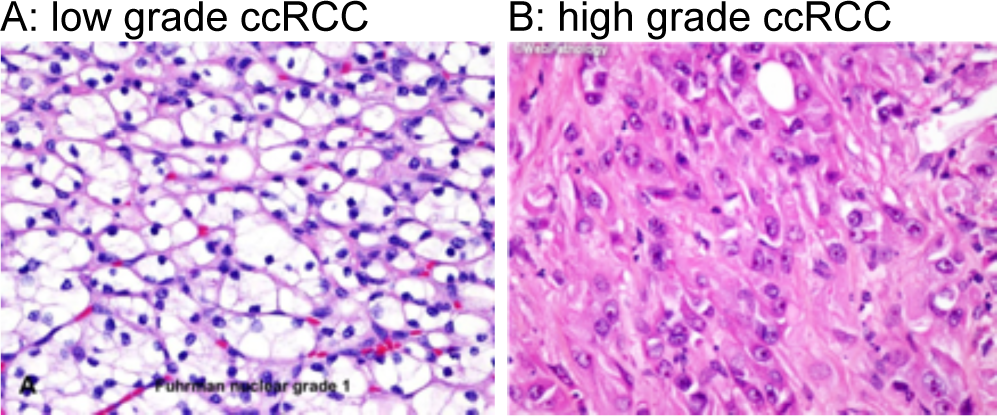
Low (A) and high (B) grade clear cell renal cell carcinoma (ccRCC), in which the stromal architectures are markedly different. Collagen/fibroblast in pink (light shaded), cancer cell nuclei in dark. Both content and distribution of fibroblasts are different between grades.

We explored several computational and mathematical modeling approaches, in order to address the following hypothesis. Within tumors, such as ccRCCs, the abundance and spatial distribution of stroma, and of CAFs in particular (“stromal architecture”), impacts tumor growth, risk of progression and response to targeted therapies. Thus, we sought to decipher the geometry and impact of cancer’s stromal architecture through the development of novel quantitative analyses. Our analyses focused on positioning of cancer-associated fibroblasts (CAFs). In the future, these analyses could be extended to the topology of other important stromal cells that often reside in the tumor microenvironment, as well as to immune cells, e.g. T-cells and macrophages^9^.

## II. Approaches and Results

CAFs shape tumor growth and therapy response, but key mechanisms of these processes are elusive: we lack quantitative methods to account for dynamical spatial distribution of CAFs and ecological interactions between CAFS and tumor^10^. Studies that integrate experimental and clinical data with mathematical modeling will inform spatial ecological processes. Such work could deepen our understanding of the impact of CAFs on tumor evolution in space and time, inform clinical choice of treatment regimens in both adjuvant and neoadjuvant settings, and offer novel angles for therapeutic interventions. To these ends, we here explored different approaches that could elucidate how spatial information of stroma and cancer cells can be analyzed, and how one could gain understanding in their temporal evolution through modeling.

### A. Geometry and distribution of cells

As a first step, we wanted to gain some understanding about the nature and impact of different geometrical arrangements between at least two different cell types that can be observed in tissues. As shown in Figure 2 (A and B), tumors might indeed evolve very different stromal architectures and resulting clustering of cells. While at this point it is unclear whether this emerges as a direct or indirect effect of CAFs themselves on tumor cells, we wanted to know how spatial measurements in the simplest settings determine the cell-to-CAF distance distributions. Cell populations surrounded by a layer of CAFs, or a regularly patterned “CAF-grid”, would result in rather different distributions (Figure 2 C and D).

**Figure 2.**
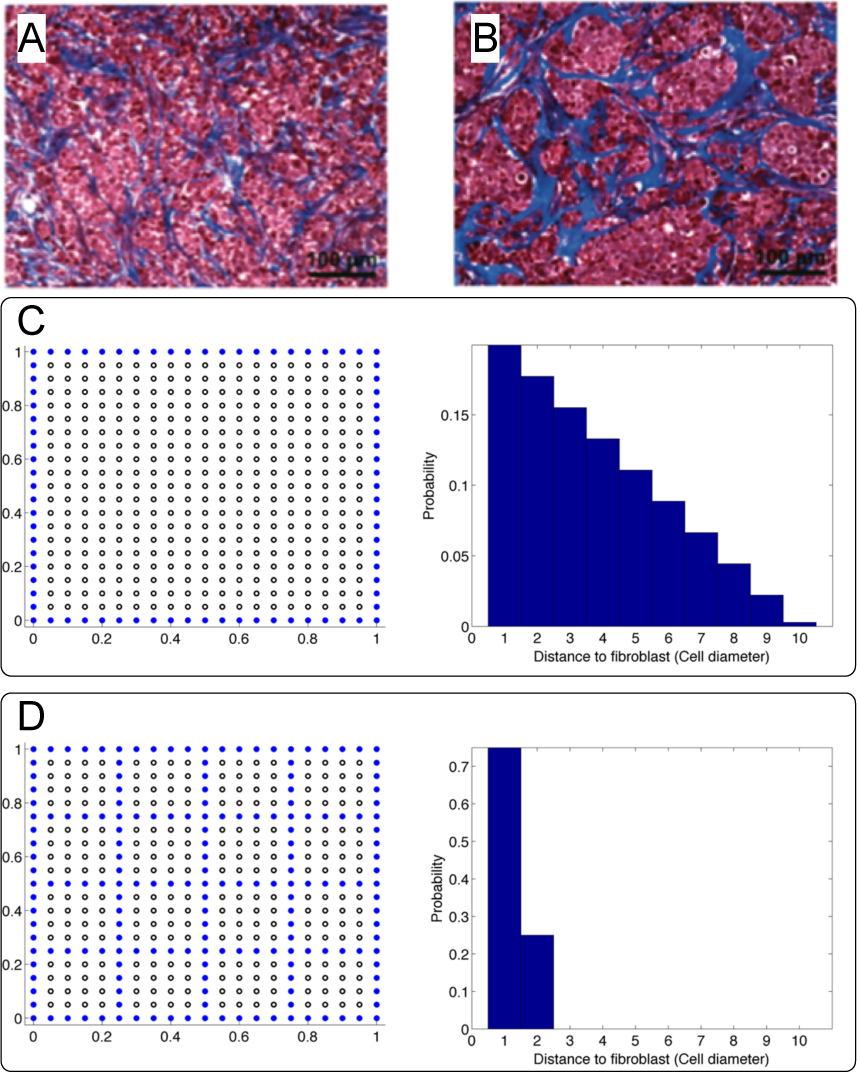
A, B: Two example images from mouse tumors from Marysyk *et al.*^11^, where tumor areas enriched in CAFs are shown in blue, and cancer cells are shown in purple. These two very different stromal geometries were shown to emerge in rather different tumors. C, D: Different geometries of CAFs lead to markedly different distributions of CAFs-cancer cell distances. Thus, if CAFs provide benefits to tumor cells during targeted treatment, these distance-statistics can have important consequences for the tumor.

Further, one could then use labeled tumor imaging information to initialize digitalized versions that could serve as the starting point for statistical, computational and mathematical analyses and predictions (Figure 3 A and B). Indeed, in an example of breast tumor tissue, proliferating cells could be found in closer proximity to CAFs (Figure 3 C and D). All together, these examples highlight the need to establish better norms of tumor image scoring that then serve to initialize predictive modeling. As the following three examples show, such modeling can be entirely computational, entirely analytical, or in hybrid form, and thus highlight different important aspects of the dynamics of non-genetically driven therapy resistance acquisition in spatially heterogeneous tumors.

**Figure 3.**
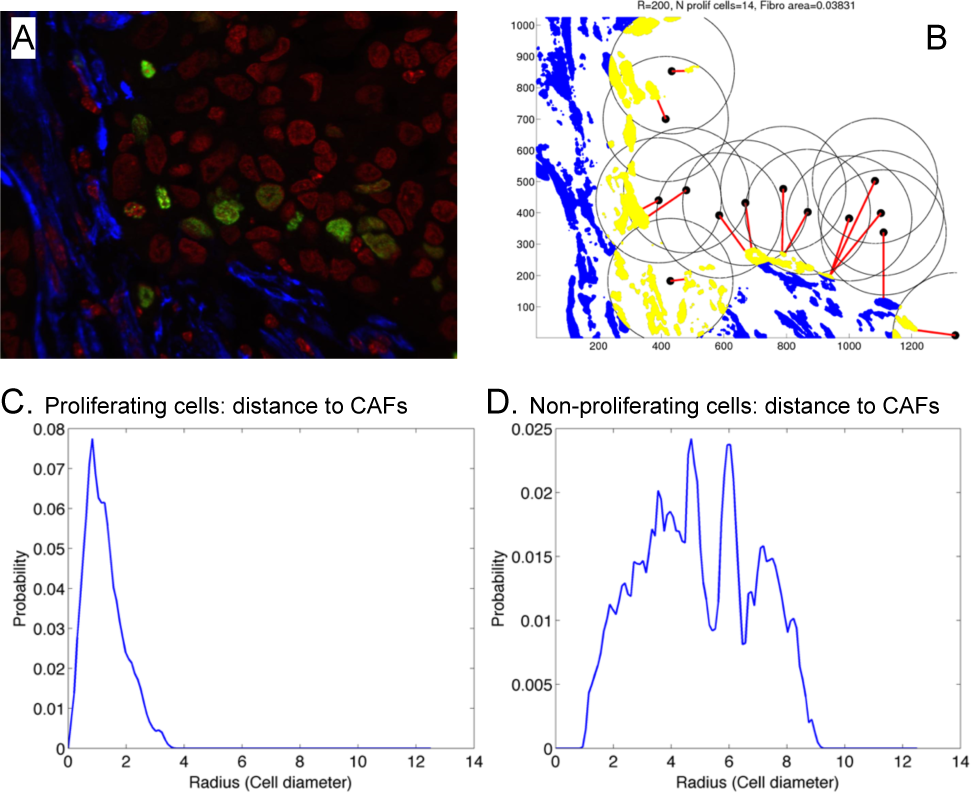
Cellular behavior in relation to stromal presence. A: Fluorescently-stained cancer tissue slide, blue: CAFs, green: proliferating cancer cells, red: non-proliferating cancer cells. B: Computational analysis of minimal distances (red), using a pre-defined radius of cancer cell-CAF interactions (gray). C: Proliferating cells’ distance to the nearest CAF. D: Non-proliferating cells’ distance to the nearest CAF.

### B. The dynamics of stromal architecture: Agent-based off-lattice approach

As a next step, we present an agent-based computational model, which has allowed us to determine the impact of fibroblast location on the evolution of pre-existing resistance to treatment in a growing or homeostatic tumor cell population. Our model contains two populations of tumor cells, labelled “sensitive” and “resistant”. Under homeostatic conditions, resistant cells have a lower growth rate than sensitive cells. Cells proliferate stochastically, with proliferation times uniformly distributed around a specified average growth rate. Under an assumption of contact-inhibition, cells may only proliferate if there is sufficient space, which (in this example) can only be made available by cell death.

The treatment we considered inhibits the proliferation of the sensitive tumor cell population, while leaving the resistant population unaffected. This confers an evolutionary advantage on the resistant cell population after treatment. However, we critically assumed that sensitive cells are able to resist the treatment in the presence of substances provided by CAFs, for example called Fibroblast Growth Factor, FGF. We here assume that FGF exists at sufficient concentrations to encourage sensitive cell proliferation only within a specified radius of a fibroblast, hence giving sensitive cells the advantage over their resistant counterparts when they are within 4 cell widths of a fibroblast.

We seeded the domain with a fixed number of fibroblasts, and varied their spatial distribution between simulation runs over time. Such initial distributions could be obtained from tumor images, as shown in Figure 4A—this will be a critical step in using agent-based modeling to simulate, test and validate evolutionary models and assumptions about CAF mediated resistance evolution. However, certain extremes of CAF distributions could also be explored: Figure 4 shows two manually generated stromal architectures as initial conditions for simulations. Our model was then used to simulate these cancer cell populations forward in time (with the CAF distribution being static). This shows that highly structured stromal architectures, e.g. in which fibroblasts are distributed in distinct clusters, permit the resistant cell population to achieve dominance much more quickly than more diverse stromal structures. Figure 4C shows the results of this time-forward modeling applied to a range of stromal architectures, indicating that random CAF distributions suppress resistant cells for longer than their more highly structured counterparts.

**Figure 4.**
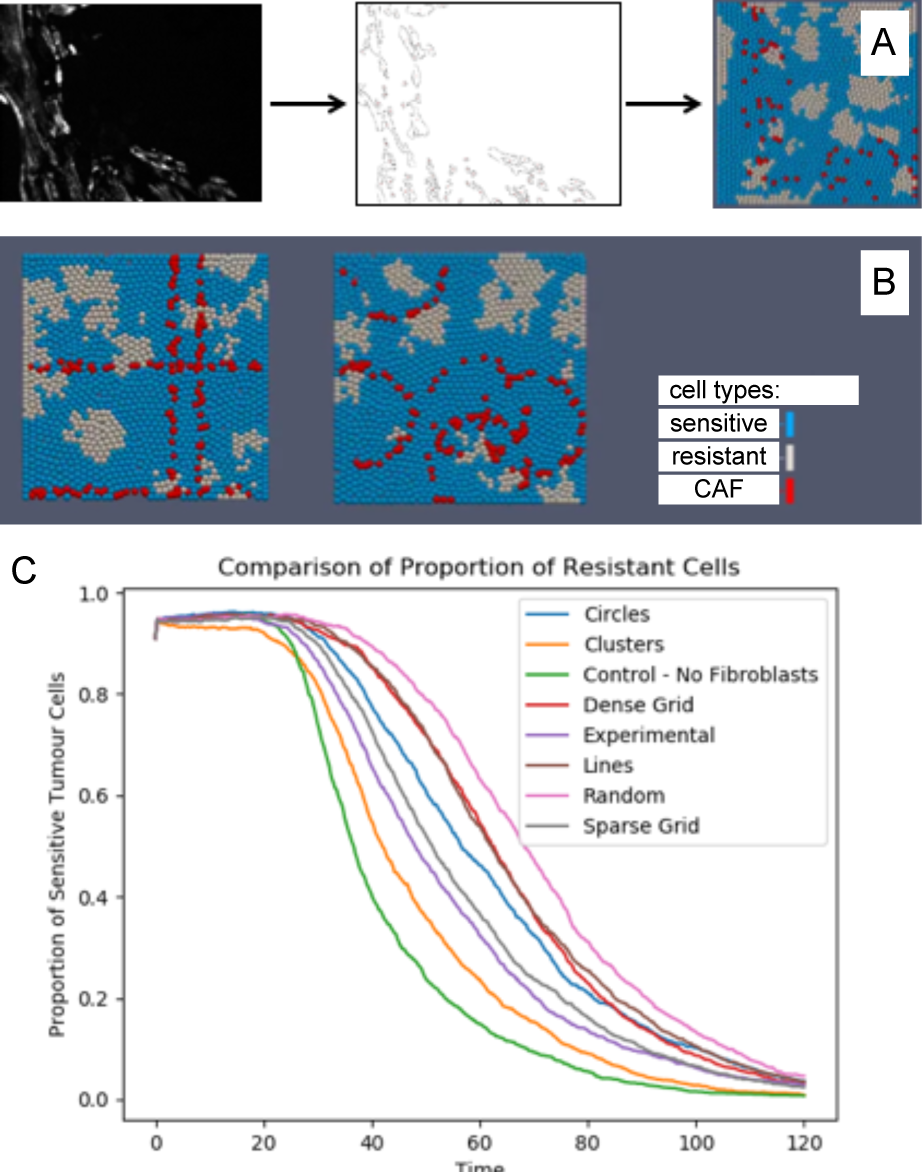
Off-lattice agent-based computation approach (using Chaste: An Open Source C++ Library for Computational Physiology and Biology^12^) in two dimensions. A: obtaining physically observed distributions of CAFs from fluorescence imaging (same as Fig. 3 A), to initialize the model. B: two examples of differing stromal architecture at the beginning of the simulation (left: “sparse grid”, right: “circles”). CAFs were assumed to evolve very slowly and not move. Treatment was “on” all the time, giving an advantage to resistant cells However, we also assumed that CAFs mitigate the selective pressure, e.g. via Fibroblast Growth Factor, at least within a radius of 4 cells. C: graphs showing the proportion of sensitive tumor cells in the population over time after treatment that started at t = 10. Resistant cells dominate the population much more quickly when stromal architecture, i.e. the CAFs, forms larger structures (e.g. circles, clusters, grids). Random CAF placement leads to slower extinction of sensitive cells.

### C. An ordinary differential equation approach to explore CAF mediated protection against targeted therapy

Tumor response to targeted therapy within a tumor is affected by the interactions between cancer cells and local microenvironments, including cancer-associated fibroblasts (CAFs). The intercellular communication between cancer and fibroblasts can be mediated by secreted growth factors or cytokines that may protect cancer cells surrounded by fibroblasts from therapy effects^13^. In order to gain valuable analytical insights, we designed an ordinary differential equation (ODE) model to describe dynamic tumor-stroma interactions in response to targeted therapy.

The tumor is classified into two subpopulations with respect to their sensitivity to targeted therapy. The tumor consists of sensitive (*S*) and resistant (*R*) cells. The stroma is implicitly in the model as a means to classify sensitivity populations further into two sub-populations. That is, we further refined the sensitive cell population as being sensitive and close to fibroblasts (*S_F_*) and sensitive but away from fibroblasts (*S_A_*). In absence of therapy, both sensitive cell populations, *S_A_* and *S_F_*, grow at the same rate (*g_s_*), while the resistant cells *R* grow at smaller rate (*g_s_ > g_R_*). We introduced a global density-dependence effect: the three populations share a carrying capacity *K*, representing the maximum packing capacity of the tumor at hand, which could be co-determined by other microenvironmental factors (such as angiogenic factors), and by the current number of dominant driver mutations (resistance mechanisms not included). By fixing *K*, we focus on short term evolution (e.g. during treatment), as clearly in the long term the whole tumour could grow further. The effect of fibroblast migration between the sensitive populations (*S_A_* and *S_F_*) is modeled by introducing transition rates α and β, with α = β (Figure 5).

**Figure 5.**
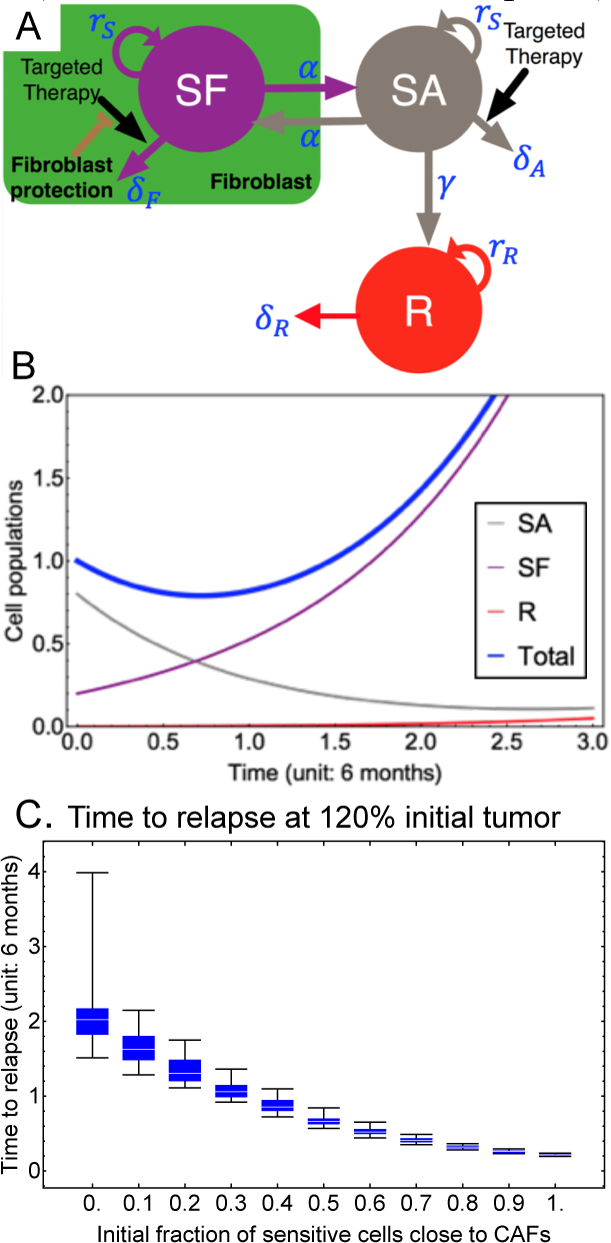
A: Schematic of the ODE two-compartment model in which therapy sensitive cells can be shielded if they are close enough to CAFs, see also Equations (1)-(3). B: simulation histories of SA (sensitive cells away from CAFs), SF (sensitive cells close to CAFs), R (resistant cells), and summation of them with the relapse in the example (blue, “Total”). Parameters: (gS,gR,K)=(1,0.5,100), (dA,dF,dR)=(2,0.05,0.05),(a=b,g)=(0.05,0.01), and (SA(0),SF(0),R(0))=(0.8,0.2,0). C: Relapse times for different initial population structures (SF(0): x-axis, SA(0)=1-SF(0), R(0)=0). For each structure, distribution of 100 simulations with 100 random transition rates (chosen from uniform distributions: aØ[0,1], gØ[0,1]) are shown by each box/whisker plot. The values for all the other parameters are same with the panel B. The time unit, 6 months, is consistent to the doubling time of renal cell carcinoma^14^.

The three populations die at rates of *δ_A_*, *δ_F_* and *δ_R_*, respectively (which could be equal in absence of therapy). We then assumed that targeted therapy induces significant cell deaths of *S_A_*. During therapy, resistance to the therapy thus emerges at a small rate, possibly due to stochastic alteration (epigenetic changes, cell-signaling changes, mutations, etc.) in the sensitive cells away from fibroblasts (*S_A_* → *R* with a rate, γ). All these dynamics can be cast into the following ordinary differential equations:

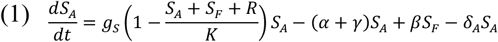

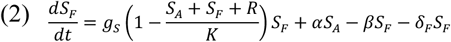

and

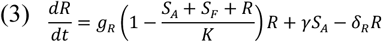

This nonlinear dynamical system is schematically drawn in Figure 5 A. Figure 5 B depicts a typical temporal response of these three populations upon targeted therapy. *S_F_* and *R* are more protected than *S_A_* under the pressure of the targeted therapy which is accounted by *δ_F,R_* < *δ_S_*. However, *R* grows in limited fashion in a short-term period due to the initial absence of resistant cells (*R*(*t* = 0) = 0) and the modeled “gcost of resistance” (*g_R_* < *g_S_*). For many choices of the model parameters, total population (or tumor size; *S_A_* + *S_F_* + *R*) decreases initially, because of the decline of *S_A_* (especially when *S_A_*(0) ≫ *S_F_*(0)). However, tumor regrows soon after due to both the fast growth of sensitive cells close to CAFs, *S_F_*(*t*) and the emergence of resistant cells *R*(*t*). The tumor eventually relapses. We focused on time to relapse following targeted therapy. In this example, we defined the time taken to reach to 120% of the initial tumor size as the relapse time. We thus observed the effect of initial proportion of *S_F_* on treatment relapse time. As expected, the more cells are protected by fibroblasts at the initial time (*S_F_*(0) ≫ *S_A_*(0), with *R*(0) = 0), the sooner the cancer relapses (Figure 5 C).

### D. Insights from a compartmentalized public goods game

Another approach that is computationally less involved than a full agent-based model, and might thus lend itself to fast dynamical forecasting of heterogeneous tumor cell-CAF populations, is the compartmentalized public goods game approach. While public goods-relationships might be seen as a way to describe intrinsic tumor cell interactions (e.g. among producers and free-riders^15^), here the public good emerges in a more complex fashion. Vascular Endothelial Growth Factor (VEGF) is provided by CAFs and drives tumor growth or protects from therapy. In addition, CAF stimulating FGF is provided by producer cells, leading to an interaction pattern in which CAFs are necessary but not sufficient providers of protection from therapy. All tumor cells are sensitive to treatment, but the more CAFs are in a cell’s vicinity, the lower the detrimental effect of treatment, which in turn is influenced by the number of producer cells. In this sense, cells close to CAF that produce stimulating factors can be labeled “resistant”—they are protected from therapy’s detrimental effects. To implicitly incorporate space into this model, we considered a finite-compartment approach^16,17^. In some compartments, cells are close to many CAFs and if they are in those compartments with few CAFs and few producer cells, the benefits of the public goods take less and less effect, or vanish entirely if no CAFs are present.

We simulated this model forward in time such that within one time-step, each compartment would experience (i) cell competitive expansion in resistant and sensitive cells during therapy, which depended on the number of CAFs present, and (ii) a random reshuffling, corresponding to cell migration and spatial heterogeneity. This dynamic is shown in Figure 6 A. Importantly, as shown in the equations in Figure 6 B, public good-mediated competition was strongest when the system was close to homeostatic carrying capacity, whereby each cell type could also proliferate and die according to their context-independent birth and death rates. In this approach, the presence of CAFs modulated the maximally achievable carrying capacity. More CAFs in one compartment would decrease the density-dependent effect implied by the carrying capacity, a form of K-selection^18^. Selection and therapy would then shift the typical number of CAFs near any sensitive cell (Figure 6 C, D).

**Figure 6.**
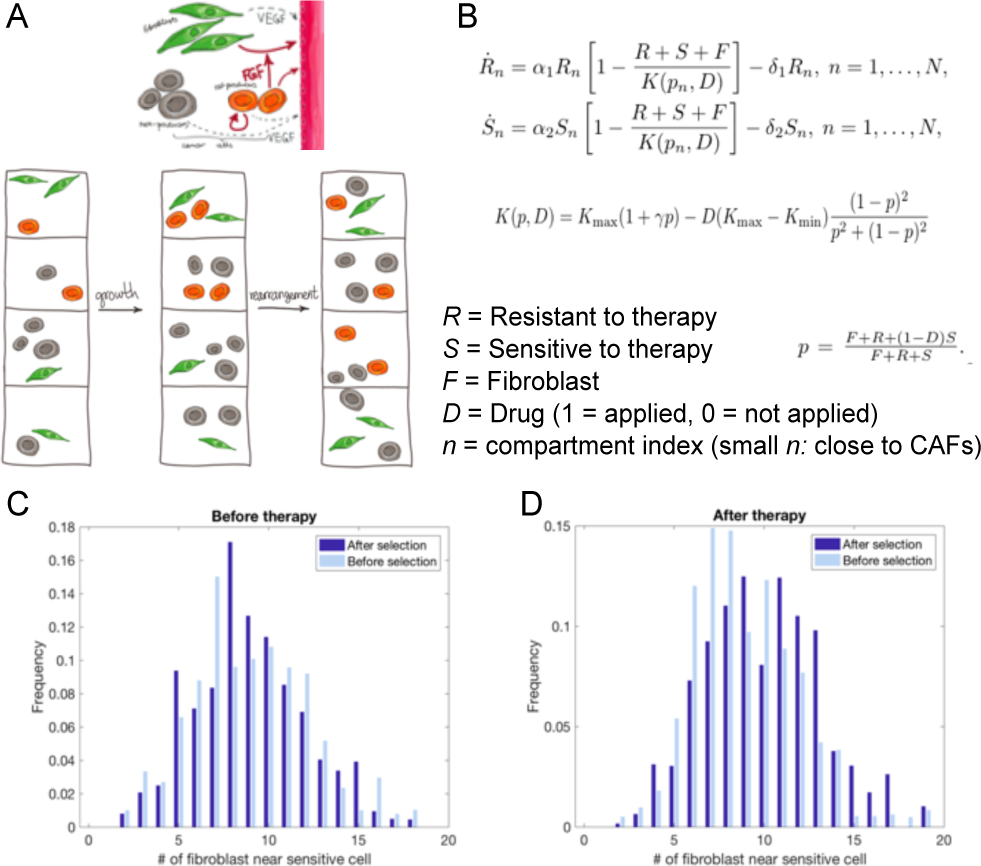
Compartmentalized public goods game approach to predict how therapy might induce higher numbers of sensitive cells near CAFs. A, top: Frequency-dependent selection and spatial reshuffling, CAFs in green, public good producer cells in red, passive free-riders in gray. A, bottom: Dynamics of selection and spatial/compartmental mixing. B: Dynamics within compartments, whereby resistant cells emerge if public good and CAFs are available. C: Distribution of CAFs near sensitive cells before and after selection, before therapy. D: Distribution of CAFs near sensitive cells before and after selection, after therapy.

## III. Summary and Outlook

We have explored how spatial heterogeneity in cancer-associated fibroblasts can affect tumor cell dynamics, and how such interactions could be modeled dynamically, in order to understand tumor-stroma co-evolution under targeted therapy.

In an off-lattice agent-based approach, we could incorporate tumor imaging of CAFs and proliferating or non-proliferating cancer cells to get a first sense of the impact of spatial stromal distributions on the extinction rate of treatment sensitive cells. This revealed that highly clustered CAF distributions might be optimal protectors of sensitive cells and subsequently act as drivers of resistance emergence and therapy failure.

Our analytical calculations have also revealed that tumors with more cells that are close to CAFs at the beginning of therapy may relapse faster. In addition, a compartment-based approach that implemented frequency-dependent selection in form of a cellular public goods game among cancer cells and between cancer cells and CAFs. This approach also showed that treatment might select for proximity to CAFs among sensitive cell populations, or stimulate CAF recruitment.

Future research should especially focus on identifying the heterogeneous spatial nature of stromal protection as observed in different cancer types and stages. On the other hand, dynamical inference from different tumor and treatment stages, but within the same patients, should be used to obtain insights into possible cellular interaction mechanisms that might be critical to gauging therapy success, or even render stromal architecture as an important co-determinant for targeted treatment administration and multi-drug considerations. Future effort should also be devoted to describe quantitatively and experimentally the connection between mechanical properties of the tumor microenvironment and the selective pressures that emerge from them^19^.

## Acknowledgment

We would like to acknowledge the Integrated Mathematical Oncology Department Chair, Dr. Alexander Anderson, for organizing the 7th Annual Moffitt IMO workshop: Stroma, where this project was conceived. We are also extremely grateful to the Moffitt Cancer Center and the Moffitt PSOC for supporting this workshop through the NCI U54CA193489 grant. BJM was supported in part by the Urology Care Foundation Research Scholar Award Program and Society for Urologic Oncology. The content is solely the responsibility of the authors and does not necessarily represent the official views of the American Urological Association (AUA) or the Urology Care Foundation.

## References

1 Spill, F., Reynolds, D. S., Kamm, R. D. & Zaman, M. H. Impact of the physical microenvironment on tumor progression and metastasis. Curr Opin Biotechnol 40, 41–48, doi:10.1016/j.copbio.2016.02.007 (2016).

2 Wilson, T. R. et al. Widespread potential for growth-factor-driven resistance to anticancer kinase inhibitors. Nature 487, 505–509, doi:10.1038/nature11249 (2012).

3 Valkenburg, K. C., de Groot, A. E. & Pienta, K. J. Targeting the tumour stroma to improve cancer therapy. Nature Reviews Clinical Oncology 15, 366–381, doi:10.1038/s41571–018–0007–1 (2018).

4 Junttila, M. R. & de Sauvage, F. J. Influence of tumour micro-environment heterogeneity on therapeutic response. Nature 501, 346–354, doi:10.1038/nature12626 (2013).

5 Hirata, E. et al. Intravital imaging reveals how BRAF inhibition generates drug-tolerant microenvironments with high integrin beta1/FAK signaling. Cancer Cell 27, 574–588, doi:10.1016/j.ccell.2015.03.008 (2015).

6 Marusyk, A. et al. Spatial Proximity to Fibroblasts Impacts Molecular Features and Therapeutic Sensitivity of Breast Cancer Cells Influencing Clinical Outcomes. Cancer Res 76, 6495–6506, doi:10.1158/0008–5472.CAN-16–1457 (2016).

7 Manley, B. J. et al. Integration of recurrent somatic mutations with clinical outcomes: a pooled analysis of 1049 patients with clear cell renal cell carcinoma. European urology focus (2016).

8 Choueiri, T. K. & Motzer, R. J. Systemic Therapy for Metastatic Renal-Cell Carcinoma. New England Journal of Medicine 376, 354–366, doi:10.1056/NEJMra1601333 (2017).

9 Seager, R. J., Hajal, C., Spill, F., Kamm, R. D. & Zaman, M. H. Dynamic interplay between tumour, stroma and immune system can drive or prevent tumour progression. Convergent science physical oncology 3, 034002, doi:10.1088/2057–1739/aa7e86 (2017).

10 Altrock, P. M., Liu, L. L. & Michor, F. The mathematics of cancer: integrating quantitative models. Nature Reviews Cancer 15, 730–745, doi:10.1038/nrc4029 (2015).

11 Marusyk, A. et al. Non-cell-autonomous driving of tumour growth supports sub-clonal heterogeneity. Nature 514, 54–58, doi:10.1038/nature13556 (2014).

12 Mirams, G. R. et al. Chaste: An Open Source C++ Library for Computational Physiology and Biology. PLOS Computational Biology 9, e1002970, doi:10.1371/journal.pcbi.1002970 (2013).

13 Heynen, G. J. J. E., Fonfara, A. & Bernards, R. Resistance to targeted cancer drugs through hepatocyte growth factor signaling. Cell Cycle 13, 3808–3817, doi:10.4161/15384101.2014.988033 (2014).

14 Nerli, R. et al. Tumor doubling time of renal cell carcinoma measured by CT. Indian J Urol 30, 153–157, doi:10.4103/0970–1591.126894 (2014).

15 Gerlee, P. & Altrock, P. M. Extinction rates in tumour public goods games. J R Soc Interface 14, doi:10.1098/rsif.2017.0342 (2017).

16 Cremer, J., Melbinger, A. & Frey, E. Growth dynamics and the evolution of cooperation in microbial populations. Sci Rep 2, 281, doi:10.1038/srep00281 (2012).

17 Kimmel, a. G., Gerlee, P., Brown, J. S. & Altrock, P. M. Neighborhood size-effects in nonlinear public goods games. bioRXiv.org 2018, https://doi.org/10.1101/347401 (2018).

18 Huang, W., Hauert, C. & Traulsen, A. Stochastic game dynamics under demographic fluctuations. Proc Natl Acad Sci U S A 112, 9064–9069, doi:10.1073/pnas.1418745112 (2015).

19 Spill, F., Bakal, C. & Mak, M. Mechanical and Systems Biology of Cancer. Computational and structural biotechnology journal 16, 237–245 (2018).

